# Agency and Embodiment Provide Dissociable Benefits to Perception, Metacognition and Neural Processing During Active Sensing

**DOI:** 10.64898/2026.10.08.757585

**Authors:** Ziying Tang, James Dowsett, Paul Taylor, Dardo N. Ferreiro, Michael Pecka

**Author notes:** These authors contributed equally (Paul Taylor, Dardo N. Ferreiro, Michael Pecka).

## Abstract

Metacognition, judging how reliable our own perceptual decisions are, is central to adaptive behavior. Such judgments hinge on the reliability of the sensory evidence on which that decision is based, and active sensing through self-directed exploration is known to improve perception itself. Yet, how agency, the authorship over self-initiated actions, and the processing of the sensory consequences, which is implicit to active sensing, modulate metacognition and its calibration, remains unknown. Whether physical embodiment adds to this beyond virtual exploration is similarly unclear, yet relevant both for daily life and for self-monitoring in agentic AI. Here, we combined a naturalistic auditory search paradigm with mobile electroencephalography. Twenty participants performed a target-detection task under three conditions: active embodied exploration, active screen-based exploration, and passive listening to replays of their own trials, while we obtained steady-state auditory evoked potentials (SSAEPs) to characterize stimulus-locked neural responses. We find that both agency conditions improved accuracy, metacognitive sensitivity, and overall SSAEP amplitude relative to passive listening, indicating that agency, rather than whole-body movement, drives these enhancements. Crucially, however, embodiment uniquely grounded both the target-evoked neural response and the subjects’ confidence to immediate sensory evidence. Thus, agency alone broadly enhances perceptual and neural processing, but embodiment specifically anchors confidence, neurally and behaviorally, to the present.

**Significance Statement:** How well we judge the reliability of our perception, called metacognitive sensitivity, matters more than ever, as we increasingly are presented with information passively through screens rather than gathering it actively via self-initiated exploration. Does actively seeking information sharpen self-monitoring, and does the medium (body vs. screen) matter? We show that active search enhances metacognitive sensitivity and brain responses alike, whether embodied or screen-based. Only embodied search, however, anchors self-evaluation to the present: Behaviorally, embodiment facilitates coupling of confidence tracking directly to current evidence rather than the previous trial’s outcome, while neuronally, it enhances responses in frontal brain regions during periods of heightened subjective confidence. This embodied grounding of self-monitoring may prove pivotal for building reliably agentic AI systems.

## Introduction

Metacognition, the capacity to evaluate one’s own cognitive processes (1), constitutes an essential foundation for adaptive behavior (2, 3). This self-monitoring is especially consequential for sensory perception: because the senses may deliver noisy and ambiguous evidence, accurately judging the reliability of a percept determines whether to commit to an action or continue gathering information. In ecological settings, sensory acquisition is rarely passive. Instead, organisms deploy active sensing through the purposive, self-directed control over when and where to sample sensory apparatuses to maximize information gain and resolve uncertainty (4–6).

During active sensing, incoming sensory streams are not arbitrary external events, but the direct consequences of self-directed exploratory actions (7–10). This closed loop between motor intent and sensory feedback gives rise to the sense of agency, i.e. the experience of authorship over one’s actions and their perceptual outcomes (11). Yet, due to the passive and open-loop design of most experimental paradigms, it is unclear how agency during sampling influences our cognition. A central question is whether agency reshapes not only what is perceived but also metacognitive calibration, i.e., how accurately the system evaluates its own perceptual decisions.

Crucially, this agency can be enacted through either embodied agency or stationary agency. The two modes differ in whether the agency is enacted by moving sensory organs through the environment (embodiment), or by a hand movement that is arbitrarily mapped onto a virtual position while the sensory organs stay relatively static (stationary). A better understanding of potential differences between these two agentic modes is increasingly significant in everyday life, where we gather more information through screens (passively scrolling feeds and actively searching alike) rather than through firsthand interaction with the physical world, raising the question of what this shift does to our metacognition and its neural basis. Specifically, agency and embodiment may not fulfill identical functional roles in shaping self-evaluation. In biological systems, for instance, perceptual accuracy and metacognitive sensitivity are dissociable, as enhanced sensory evidence can inflate subjective confidence without improving its calibration to objective accuracy (2, 12). Agency could plausibly improve metacognitive sensitivity by supplying the metacognitive system with self-generated predictive signals that iteratively track evidence reliability (10, 13, 14). Conversely, embodiment might provide a distinct, complementary contribution via body-anchored sensorimotor contingencies that ground confidence directly in organism–environment coupling (8, 15). Whether agency and embodiment make dissociable contributions to metacognitive monitoring remains an open question.

Beyond our daily life, the same distinction aligns with contemporary debates in artificial intelligence (AI) regarding the role of embodiment in generating autonomous agency (16, 17). As AI advances toward agentic architectures, enabling calibrated self-monitoring remains a key challenge (18). Recent syntheses indicate that, although substantial progress has been made in developing autonomous systems, current architectures still face challenges in supporting adaptability and in translating perceptual intelligence into effective behavior (19, 20). Emerging evidence suggests that embodiment, through closed-loop sensorimotor interaction with the environment, may be essential (21), and it likewise shapes whether observers attribute agency to an artificial system at all (22). Understanding how biological systems combine or dissociate agency and embodiment to refine self-evaluation could thus provide foundational architectural principles for building more reliable autonomous agents.

Active sensing confers documented perceptual advantages across sensory modalities by aligning neural excitability with the temporal structure of incoming stimulation (23, 24). In the auditory domain, self-directed exploration enables listeners to control the timing and sequence of sensory sampling, generating efference copies that sharpen sensory predictions and reduce perceptual uncertainty (13, 25). In its fully embodied form, active sensing couples the organism to its environment, modulating neural response during exploration (26–29). To capture this interactive complexity, the Sensory Island Task (SIT), originally developed for freely moving non-human animals (30) and subsequently adapted for human participants (31), provides an ecologically valid framework for investigating active auditory perception by embedding participants in an intuitive, naturalistic task. Notably, we previously established that auditory frequency discrimination during embodied exploration in SITh matches performance in stationary settings (31, 32). Whether this parity in first-order performance and perception accuracy extends to second-order, metacognitive sensitivity, however, remains unknown.

If active sensing supplies the metacognitive system with better information about the state of the perceptual system, then it could also sharpen metacognitive sensitivity, i.e., how well confidence tracks accuracy. Crucially, a stronger neural response to the stimulus does not guarantee a sharper correspondence between confidence and accuracy (2), because amplifying the sensory signal can inflate confidence overall without improving how well it discriminates correct from incorrect judgments (33). A genuine improvement in metacognition should therefore appear as confidence that more sharply separates correct from incorrect judgments, without any overall rise in confidence.

To investigate the neural processing of auditory perception and metacognitive monitoring within an ecological framework, we implemented mobile EEG recordings during SITh, which captures the rapid ongoing details of unrestricted behavior and the neural activity during task performance (34, 35). We collected steady-state auditory evoked potentials (SSAEPs), also known as auditory steady-state response (ASSR), to quantify the extent to which repetitive auditory stimulation elicits phase-locked cortical activity (36, 37). Their high spectral signal-to-noise ratio renders them particularly suitable for artifact-prone environments, including mobile EEG recording during locomotion (27, 38). We use SSAEPs as a neural readout of stimulus processing during naturalistic sensory search, thereby establishing a methodological framework for integrating mobile neural recordings with metacognitive research.

Specifically, this study isolates the two components: effects driven by agency should appear equally under embodied and screen-based active search, whereas effects that require embodiment should appear under whole-body search alone. We find that, first, agency improves both perceptual accuracy and metacognitive sensitivity relative to passive listening, and that this benefit stems from self-directed control, instead of whole-body movement per se. Second, agency broadly amplifies the SSAEP amplitude compared to passive listening of the same stimuli. Critically, only under embodied exploration was this enhanced neural response sensitive to confidence: target-evoked amplitude was larger on high-than on low-confidence trials, while the response to non-target sounds was unaffected, and self-evaluation remained anchored to the present sensory evidence. These effects are absent during screen-based search despite matched behavior. Together, these findings indicate that embodiment, in particular, shapes not only what is perceived but how accurately metacognition tracks the reliability of concurrent auditory perception.

## Results

We adapted the SITh paradigm (31) into three conditions to test the influence of agency and embodiment on auditory perception: embodied agency (EA), self-directed search through whole-body locomotion; stationary agency (SA), self-directed screen-based stationary exploration; and non-agency (NA), passively listening to the sound sequences heard during the SA condition (Fig. 1). Continuous mobile EEG was recorded during the whole process. On each trial, participants searched for a target sound island (500 Hz) concealed among three non-target islands (460–540 Hz; Fig. 1A). Islands were invisible and had randomized locations (SI Appendix, Figs. S1 and S2), while a non-island stimulus (850 Hz) was presented outside all islands (Fig. 1A). All stimuli consisted of 50 ms pure tone sound pips and were delivered at a fixed repetition rate of 4 Hz to elicit SSAEPs. Upon identifying the putative target island, participants pressed a button and proceeded to rate their confidence on a scale from 1 (completely uncertain) to 10 (completely certain; Fig. 1B). We first validated the mobile SSAEP recordings, then examined how agency and embodiment shaped perceptual performance, metacognitive sensitivity, and brain response.

**Figure 1:**
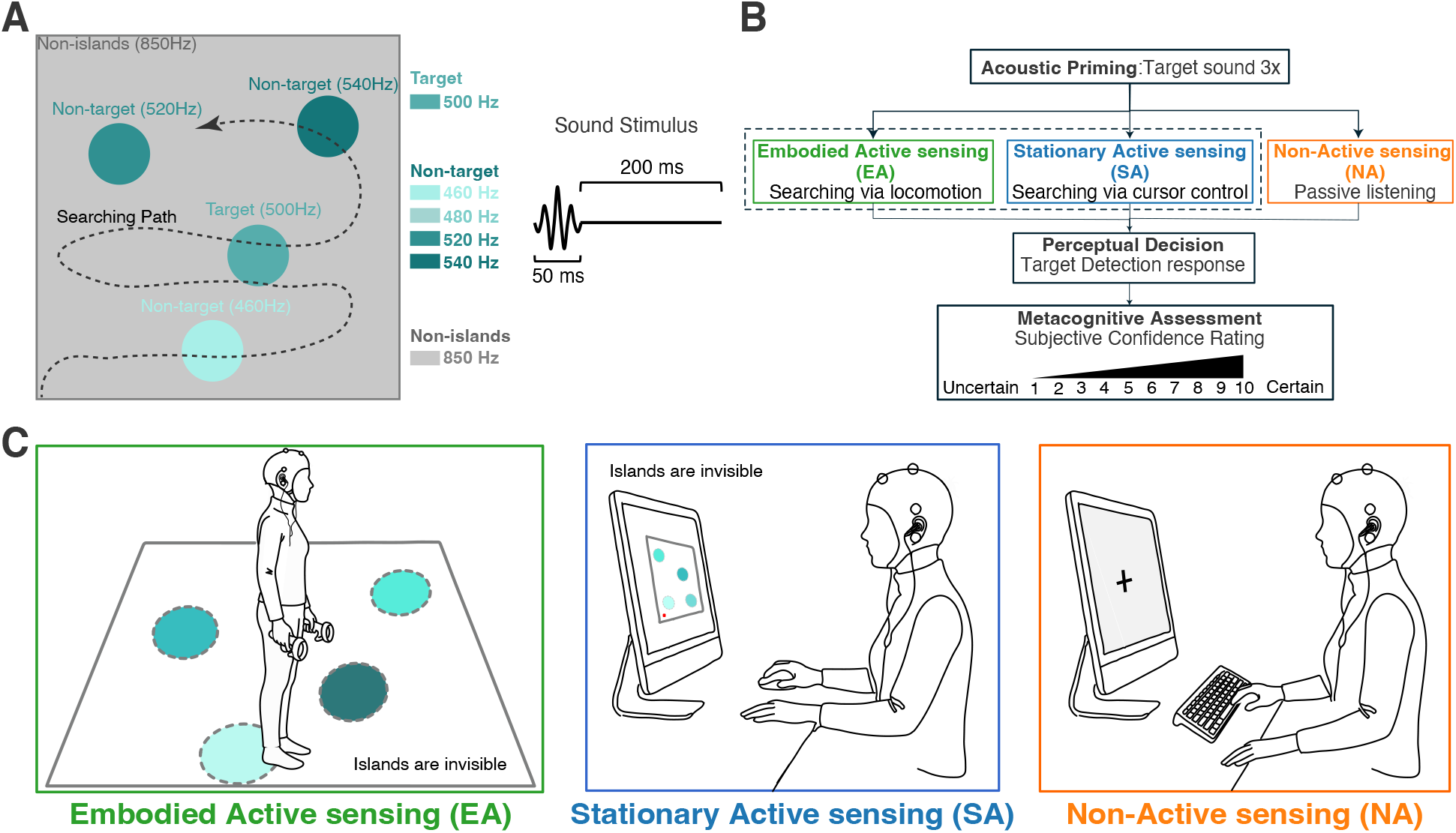
Multi-islands Sensory Island Task for Humans (SITh). Participants search an arena for a target sound island, using sound frequency as the only cue. (**A**) The square arena contains four islands—one target (500 Hz) and three non-targets (drawn from 460, 480, 520, 540 Hz) —embedded in non-island space (850 Hz). Sound stimuli are triggered dynamically from the real-time position of the participant (Embodied Agency, EA) or cursor (Stationary Agency, SA) and presented at a fixed rate of 4 Hz (50-ms tone + 200-ms inter-stimulus interval). (**B**) Each trial opens with three target-sound repetitions, followed by active search (EA, SA) or passive replay of a matched soundtrack (Non-Agency, NA). Participants press a button upon detecting the putative target sound, then rate confidence on a 1–10 scale. (**C**) The three conditions: EA, search by embodied movement in a 3 *×* 3 m arena; SA, search by moving a virtual cursor (red spot) in a 10 *×* 10 cm on-screen arena; NA, stationary passive listening to matched playback while fixating a central cross. Islands in EA and SA are drawn for illustration only and are invisible to participants.

### Feasibility and Validation of mobile SSAEPs

Spectral and reliability analyses demonstrated that 4 Hz SSAEPs serve as a robust, artifact-resistant measure of neural processing across both stationary and locomotive conditions. Normalized power spectral density (PSD) profiles exhibited distinct, sharp peaks at the 4 Hz fundamental frequency and its harmonics (8, 12, 16, and 20 Hz), with no comparable activity detected during inter-trial intervals (Fig. 2A). To assess the temporal stability of this signal at the frontal midline electrode (Fz), we computed test-retest reliability via a half-split waveform comparison (detailed further in Methods); reliability for the non-island stimulus remained consistently high and statistically indistinguishable across varying locomotion demands (EA: *r* = 0.91*±*0.09; SA: *r* = 0.90*±*0.11; NA: *r* = 0.86*±*0.12; Friedman test, *χ*^2^(2) = 0.70, *p* = .70; Fig. 2B), with individual consistency coefficients ranging from 0.57 to 0.99. In every condition, group-level reliability far exceeded a simulated null distribution of random half-waveform correlations (all *p* < .001, Bonferroni-corrected across conditions), confirming that the response recorded at Fz was reproducible within and across participants well above chance. Control epochs analyzed at a non-harmonic and non-stimulus frequency (3.1 Hz) produced near-flat time-domain waveforms, contrasting with the structured, stimulus-specific responses observed at 4 Hz (Fig. 2C); this contrast forms the basis of our primary neural measure, the normalized amplitude, the SSAEP peak-to-trough amplitude of a given stimulus relative to that of control epochs (detailed further in Methods). Furthermore, the waveforms from other channels are provided in SI Appendix, Fig. S3. Collectively, these results confirm that the recorded signals represent authentic, stimulus-locked neural responses rather than motion artifacts or spectral leakage and support the feasibility of employing SSAEPs during unrestricted embodied exploration.

**Figure 2:**
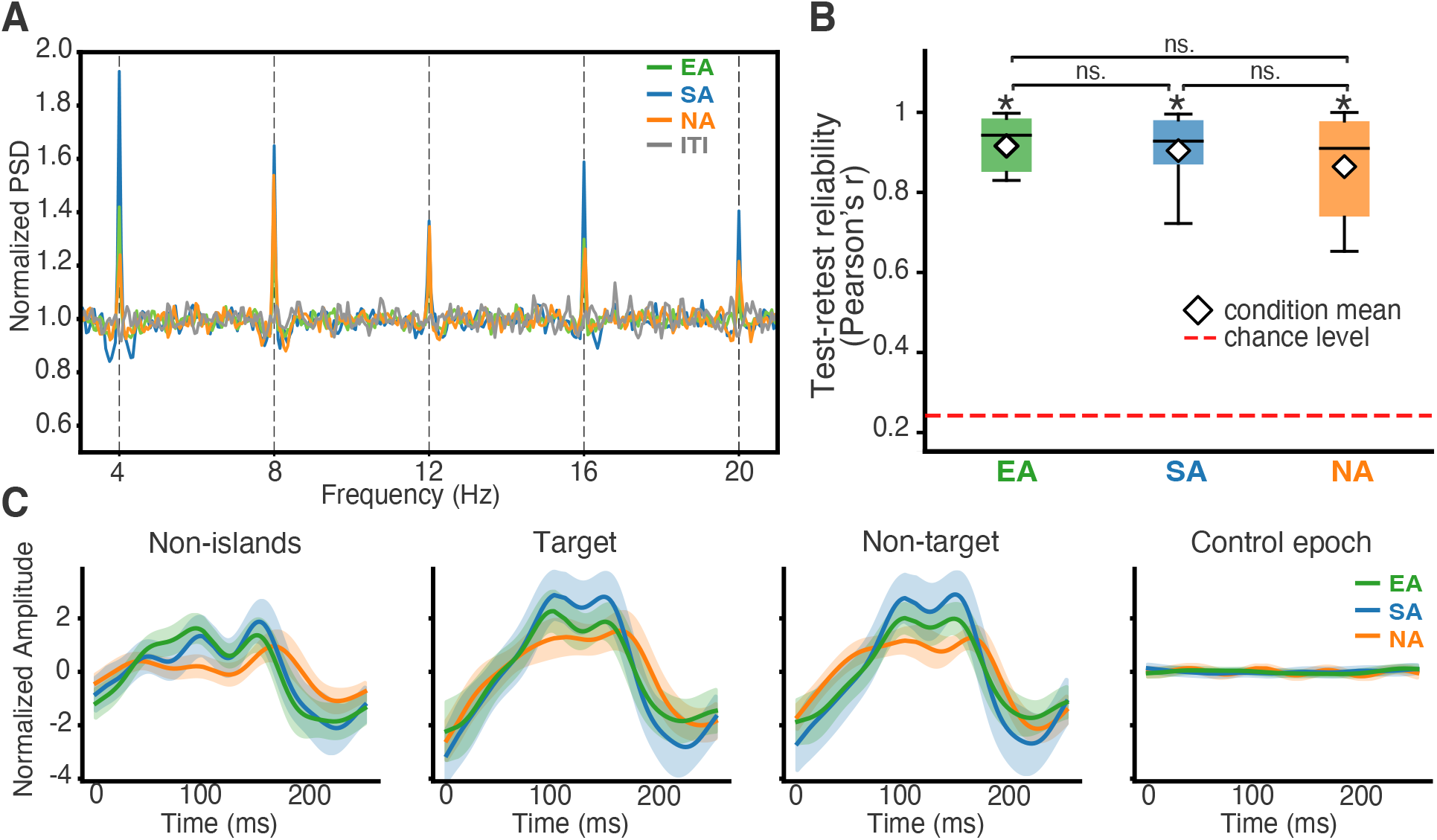
Methodological Validation and Reliability of Fz-channel SSAEPs. (**A**) Normalized power spectral density (PSD) during auditory stimulation. Spectra show robust peaks at 4 Hz and its harmonics (8, 12, 16, 20 Hz) across the EA, SA, and NA conditions during sound-on periods, with no such peaks during sound-off periods (inter-trial intervals, ITIs). (**B**) Test–retest reliability of SSAEPs. Boxplots show split-half Pearson correlations between non-island waveforms at Fz. The red dashed line indicates the mean of a spectrally matched chance level generated by Fourier phase randomization (EA = 0.249, SA = 0.238, NA = 0.228). All conditions show high temporal stability and consistency of the evoked responses (EA: *t*(19) = 43.04, *p* < .001; SA: *t*(19) = 36.85, *p* < .001; NA: *t*(19) = 31.13, *p* < .001, Bonferroni corrected). Reliability did not differ significantly across conditions (Friedman *χ*^2^(2) = 0.70, *p* = .70). (**C**) Evoked waveforms across sound stimuli. Time-domain SSAEP waveforms for non-island, target, and non-target sounds exhibit distinct, component-specific morphologies, in contrast to the near-flat response obtained from control epochs analyzed at the 3.1 Hz non-stimulus frequency. Waveforms span a full 250 ms cycle (0–250 ms), with shaded error bands.

### Active Exploration Enhances Perceptual Accuracy and Metacognitive Sensitivity

Active auditory exploration yielded greater perceptual accuracy and metacognitive sensitivity than passive listening, regardless of whether embodied movement was involved (Fig. 3). Perceptual accuracy showed a significant main effect of condition (repeated-measures ANOVA, *F*(2,38) = 10.00, *p* < .001; Fig. 3A), with both embodied agency (EA) and stationary agency (SA) outperforming non-agency (NA) (all *p* < .01, FDR-corrected) and no difference between the two agency conditions (*p* = .80; Fig. 3A). Subjective normalized confidence ratings, by contrast, remained equivalent across all conditions (repeated-measures ANOVA, *F*(2,38) = 1.57, *p* = .22; Fig. 3B; the same holds for raw average confidence, *F*(2,38) = 0.32, *p* = .73; SI Appendix, Fig. S4), indicating that perceived performance was well-matched from the participants’ perspective. We quantified metacognitive sensitivity as the area under the type-2 receiver operating characteristic curve (AUROC2), which captures how well a participant’s confidence discriminates correct from incorrect judgments — values near 0.5 indicate confidence unrelated to accuracy, whereas values approaching 1 indicate near-perfect discrimination (39, 40). Metacognitive sensitivity followed the same pattern as accuracy, with a significant main effect of condition (*F*(2,38) = 10.25, *p* < .001; Fig. 3C), both agency conditions exceeding passive listening (all *p* < .01, FDR-corrected), and no difference between them (*p* = .62; Fig. 3C); the type-2 ROC curves of a representative participant illustrate this advantage (Fig. 3D). Confirming that confidence carried genuine metacognitive information, it was reliably higher on correct than incorrect trials in every condition (SI Appendix, Fig. S6). Because subjective confidence was equivalent across conditions, this AUROC2 advantage cannot be attributed to shifts in subjective task demands, indicating that agency, rather than embodied movement, is the primary driver of enhanced perceptual and metacognitive performance. In addition, participants’ musical ability was positively associated with AUROC2 across all conditions (Figs. 3E, F).

**Figure 3:**
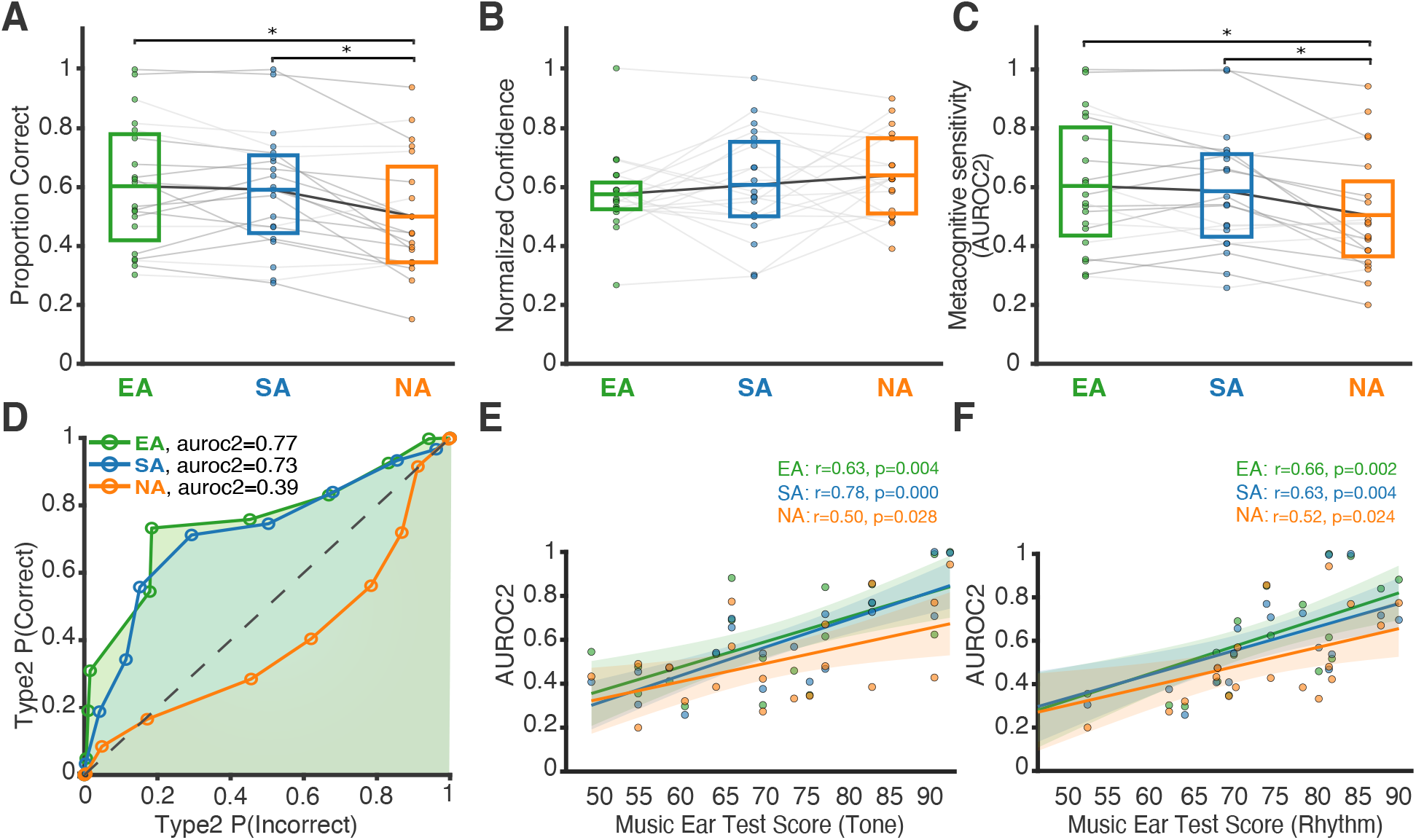
Improved Behavioral Performance and Metacognitive Sensitivity under Active Exploration. (**A**) Perceptual Accuracy. Accuracy boost within agency (EA, SA) compared to passive listening (NA, all *p* < .01). (**B**) Normalized Confidence Ratings. Mean normalized subjective confidence by condition; no significant modulation was found across the three conditions (*p* = .22), which ensures that shifts in subjective perceived performance or response bias do not confound AUROC2 values. (**C**) Metacognitive Sensitivity. Group-level values confirm significantly improved performance monitoring within the agency (EA and SA, all *p* < .01). (**D**) AUROC2 from Type-2 ROC analysis. Type-2 ROC curves from one participant across the three conditions. The area under the curve is AUROC2. Individual participant curves are shown separately in SI Appendix, Fig. S5. (**E, F**) Positive correlation between musical competence (MET) and metacognitive sensitivity (AUROC_2_). Melodic (Tone; **E**) and rhythmic (Rhythm; **F**) MET scores were both significantly and positively correlated with AUROC2 in every condition (EA, SA, NA; all *p* < .05, FDR-corrected), indicating a stable link between auditory proficiency and metacognitive sensitivity that persists independently of task condition. Individual data points are shown with regression lines and 95% confidence intervals (shaded areas); Pearson r and p-values are given in each plot legend.

Beyond these condition-level effects, we examined trial-by-trial confidence dynamics to test whether self-evaluation depends on trial history (Fig. 4). Within each condition we compared current confidence following high-versus low-confidence trials, and following correct versus incorrect trials, using Wilcoxon signed-rank tests. Current confidence showed clear hysteresis with respect to prior confidence: it was significantly higher after high-than after low-confidence trials in both agency conditions (EA: *N* = 18, *p* < .05, Fig. 4A; SA: *N* = 19, *p* = .03, Fig. 4B), but not during passive listening (NA: *N* = 20, *p* = .63, Fig. 4C). The influence of prior objective performance was selective (participants received feedback about correctness after each trial): confidence under embodied agency was independent of prior accuracy (EA: *N* = 19, *p* = .23, Fig. 4D), whereas under stationary agency a correct preceding trial significantly increased current confidence (SA: *N* = 19, *p* < .001, Fig. 4E); passive listening likewise showed no influence of prior accuracy (NA: *N* = 20, *p* = .77, Fig. 4F). Thus, serial dependence on prior confidence persisted across both active modes, but prior performance modulated self-evaluation exclusively under stationary agency.

**Figure 4:**
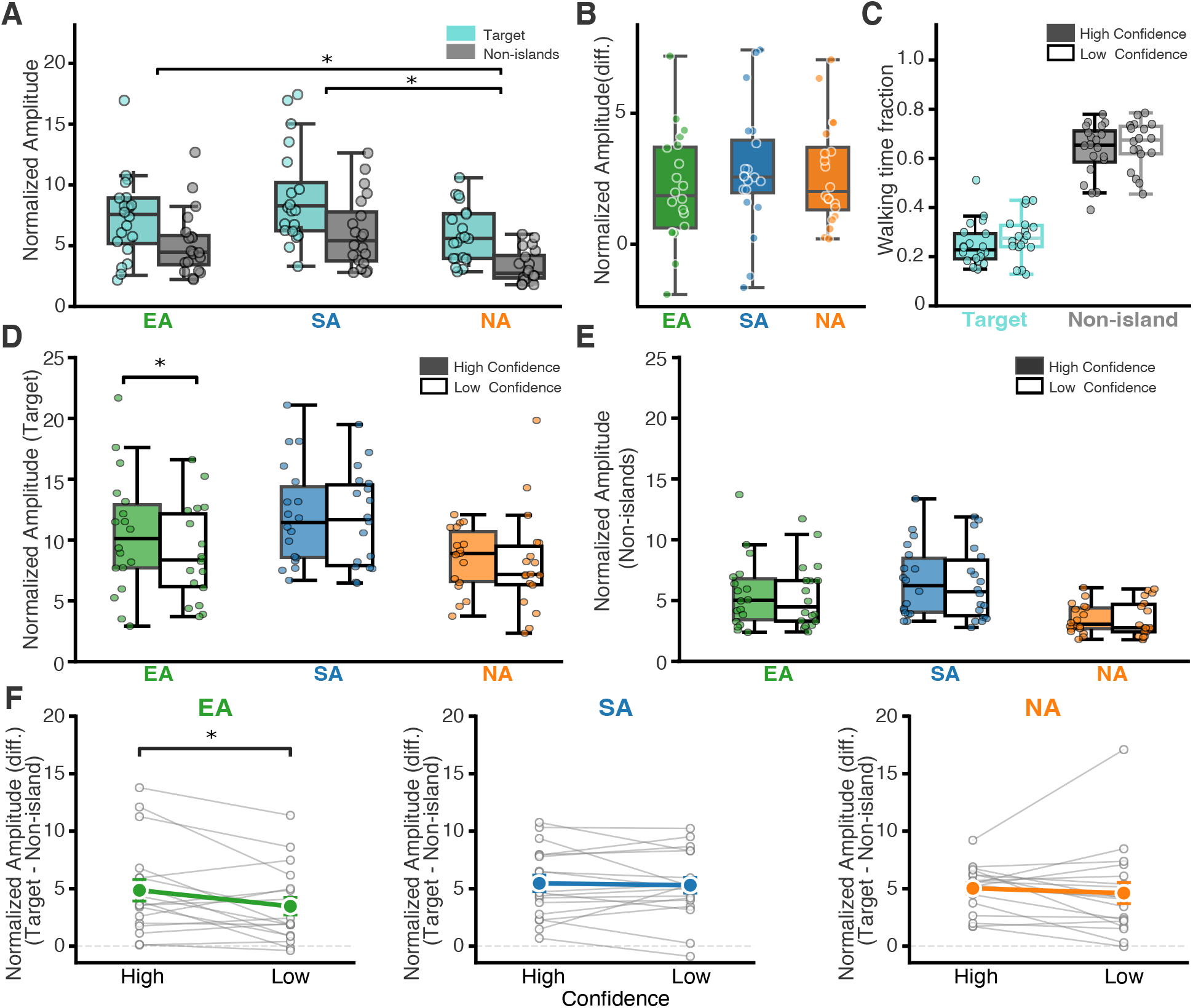
Prior Confidence and Performance Differentially Modulate Current Confidence across Agency Conditions. Current-trial confidence split by the preceding trial’s confidence status (A, B, C) or the preceding trial’s accuracy (D, E, F), shown separately for EA, SA, and NA; each panel shows the normalized confidence distribution with a participant-level mean boxplot. Within-condition comparisons used Wilcoxon signed-rank tests. (A, B, C) Prior confidence: high prior confidence significantly elevated current confidence under both EA (*N* = 18, *p* < .05; A) and SA (*N* = 19, *p* = .03; B), but not NA (*N* = 20, *p* = .63; C). (D, E, F) Prior performance: a correct preceding trial significantly increased current confidence under SA (*N* = 19, *p* < .001; E), but not under EA (*N* = 19, *p* = .23; D) or NA (*N* = 20, *p* = .77; F). N differs across panels because participants without low-confidence trials or incorrect trials could not enter the corresponding split. Boxplots display medians, interquartile ranges, and individual participant data points.

### Active Exploration Enhances SSAEP Amplitude, and the Target Response is Sensitive to Confidence under Embodied Agency

Finally, we determined to what extent the effects of exploration type were also reflected in EEG recordings. Our analysis revealed that agency uniformly increases normalized amplitude at the frontocentral electrode (Fz) for both target and non-island responses (Fig. 5). A 3 (condition) *×* 2 (stimulus) repeated-measures analysis, performed with the aligned rank transform (ART) revealed significant main effects of condition (*F*(2,38) = 10.59, *p* < .001; Fig. 5A) and stimulus (*F*(1,19) = 57.26, *p* < .001; Fig. 5A), and no condition *×* stimulus interaction (*F*(2,38) = 0.10, *p* = .90; Fig. 5B). Pairwise comparisons of the condition marginal means (justified by the absent interaction) showed higher normalized amplitude in both agency conditions than in passive listening (EA vs NA, *p* < .05, *d* = 0.74; SA vs NA, *p* < .001, *d* = 1.07). The difference between the two agency conditions did not reach significance, though there was a trend toward higher amplitude in SA than EA (*p* = .077, *d* = 0.42). Thus, agency broadly increased SSAEP amplitude, while the target/non-island selectivity remained stable across conditions (Fig. 5B). Within this stable response structure, we asked whether the target-evoked response was sensitive to confidence (*N* = 18; participants 7 and 17 lacked low-confidence trials). Target-normalized amplitude at Fz was higher on high-than on low-confidence trials (main effect of confidence: *F*(1,17) = 12.67, *p* = .002); in post hoc comparisons this difference was significant only under EA (*p* = .03, FDR-corrected; Fig. 5D) but not under SA or NA (condition *×* confidence interaction did not reach significance, *F*(2,34) = 1.29, *p* = .29). By contrast, the non-island response was not sensitive to confidence in any condition (*F*(1,17) = 1.57, *p* = .23; Fig. 5E), and neither were the non-target islands (SI Appendix, Fig. S7B). Furthermore, the fraction of time spent walking during stimulus presentation did not differ between high- and low-confidence trials (Fig. 5C; see SI Appendix, Fig. S7A for all stimulus types), indicating that the confidence-related difference was not a byproduct of differential movement, and rules out walking-induced motion artifacts or movement-related arousal as its source. To test whether confidence modulated target processing specifically, we entered stimulus as a factor in a 3 (condition) *×* 2 *×* (stimulus) 2 (confidence) repeated-measures analysis. This analysis showed that confidence interacted with stimulus (*F*(1,17) = 11.24, *p* = .004), with the selectivity margin being larger on high-than on low-confidence trials. Neither the condition *×* stimulus interaction (*F*(2,34) = 2.04, *p* = .15) nor the three-way interaction (*F*(2,34) = 1.82, *p* = .18) reached significance, indicating that basic stimulus selectivity was preserved across conditions and that the confidence-related shift in selectivity was not statistically distinguishable between them. In condition-wise comparisons, however, the margin shifted with confidence only under embodied agency (*t*(17) = 2.38, *p* = .03, *d* = 0.56), and not under stationary agency (*p* = .72) or passive listening (*p* = .54, Fig. 5F). Confidence therefore modulated target processing specifically, and did so most clearly during embodied search. Yet this confidence-related difference did not reflect accuracy: target-evoked amplitude in the active conditions did not differ between correct and incorrect trials, and a correct–incorrect difference emerged only during passive listening (SI Appendix, Fig. S8). Together, these results show that agency broadly amplifies the SSAEP response to sounds, while the target-evoked response is sensitive to confidence, most clearly during embodied exploration, a difference specific to the sensory target and not explained by differences in movement.

**Figure 5:**
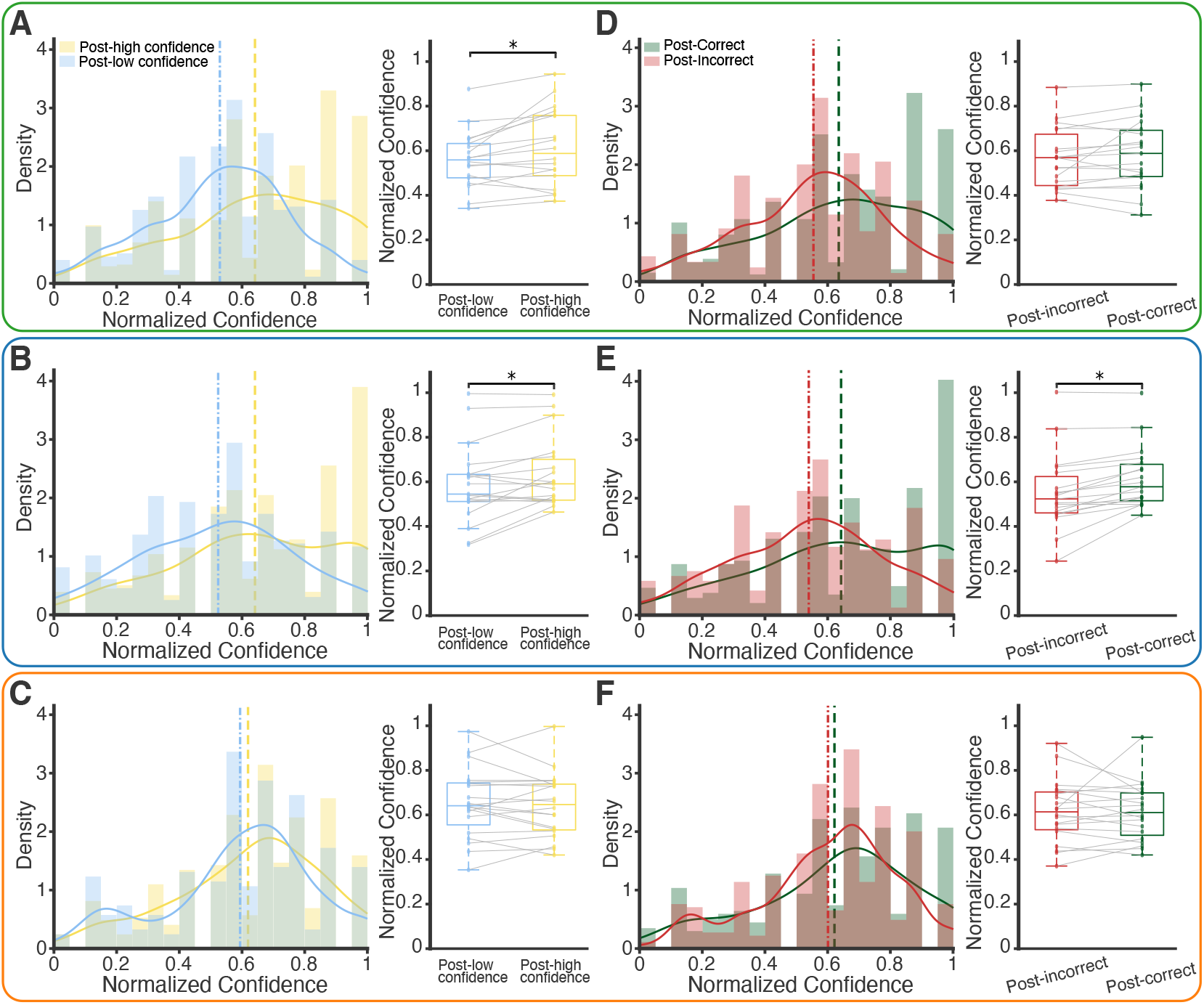
SSAEP Amplitude at Electrode Fz and Its Sensitivity to Confidence. Normalized SSAEP peak-to-trough amplitude at electrode Fz. Condition and stimulus effects (A, B) were tested with a 3 (condition: EA, SA, NA) × 2 (stimulus: target, non-island) aligned rank transform (ART) repeated-measures ANOVA (*N* = 20); confidence-dependent analyses (C–F) contrast high-versus low-confidence trials (*N* = 18; participants 7 and 17 excluded for lacking low-confidence trials). (**A**) Agency increases normalized SSAEP amplitude at Fz. Normalized amplitudes for target and non-island stimuli across conditions. Both main effects were significant, condition (*F*(2,38) = 10.59, *p* < .001) and stimulus (*F*(1,19) = 57.26, *p* < .001), and pairwise comparisons of the condition marginal means showed higher amplitude in both agency conditions than in passive listening (EA vs NA, *p* < .05; SA vs NA, *p* < .001), with no difference between the two agency conditions (EA vs SA, *p* = .077). (**B**) Invariance of stimulus-driven selectivity. The selectivity margin (target minus non-island normalized amplitude) did not differ across conditions (condition *×* stimulus interaction: *F*(2,38) = 0.10, *p* = .90), indicating that basic sensory selectivity is preserved regardless of agency or passive listening. (**C**) Invariance of locomotion across confidence states during EA. Fraction of time spent walking during stimulus presentation for the target and non-island stimuli in the EA condition, derived from accelerometer data and split by confidence state. Walking duration did not differ significantly between high- and low-confidence trials for either the target (*p* = .35) or the non-island stimulus (*p* = .99; *N* = 18, paired t-tests), ruling out walking-induced motion artifacts or movement-related arousal as the driver of the confidence-related difference in target SSAEP amplitude (Fig. 5D). (**D**) Target amplitude is sensitive to confidence under EA. Target-evoked SSAEP amplitudes as a function of confidence state. A significant enhancement in target amplitude during high versus low confidence states emerged exclusively during EA (*p* = .03). (**E**) Stability of non-island processing across confidence states. Non-island amplitude showed no significant modulation by confidence level in any condition (no effect of confidence, *F*(1,17) = 1.57, *p* = .23). Boxplots display medians, interquartile ranges, and individual subject data points. (**F**) Confidence modulates target processing specifically. Selectivity margin (target minus non-island normalized amplitude) at Fz by confidence state. Gray lines, individual participants; colored points, mean ±SEM. The stimulus *×* confidence interaction was significant (3 *×* 2 *×* 2 ART ANOVA: *F*(1,17) = 11.24, *p* = .004); the three-way interaction was not (*F*(2,34) = 1.82, *p* = .18). The margin shifted with confidence only under EA (*p* = .03).

## Discussion

Agency and embodiment make dissociable contributions to metacognition. Agency, whether embodied or screen-based, enhanced perceptual accuracy, metacognitive sensitivity, and the overall neural response while preserving the target/non-island distinction. In addition, embodiment made a further, separable contribution: confidence was insulated from prior performance, and the target-evoked response tracked current confidence most clearly, with the target-over-non-island margin widening on high-confidence trials.

Both agency-driven conditions significantly enhanced perceptual accuracy, implying that agency, rather than whole-body movement per se, is the key variable conferring these perceptual advantages. Because incidental physical effort has not been found to impair metacognitive accuracy (41), this behavioral equivalence more likely reflects shared control over sensory sampling than an offset of motor costs. Active sensing could plausibly either generate self-structured sensorimotor flow in which behaviorally relevant invariants, such as acoustic affordances, become more directly detectable (8, 42), or it may supply efference copies that sharpen internal sensory predictions and reduce perceptual uncertainty (10, 13, 14). In addition, metacognitive sensitivity also scaled with individual musical ability across all conditions, indicating a stable perceptual baseline that agency adds to rather than creates, consistent with domain-specific accounts in which metacognitive sensitivity is inherited from the fidelity of first-order perceptual representations (12, 43, 44).

Building on this picture of agency-driven improvements in perceptual accuracy, the enhancement of metacognitive sensitivity points to a deeper functional role for agency in the action-perception cycle. Metacognition may not only be a post-decisional commentary on an already-formed percept, but an ongoing inferential process (3, 45, 46), a functional gate determining whether the perceptual cycle closes in commitment or continues through further sampling (47). This gate is only fully operative when action is available: passive listeners cannot restructure the sensory evidence in response to low confidence, whereas agency restores the loop that ties confidence to perceptual reliability, consistent with the meta-listening framework, in which metacognition depends on the precision of internal predictions, heightened when listeners control the flow of evidence (48). Agency thus completes the action–perception–metacognition loop.

Consistent with this closed loop, confidence on each trial was not formed in isolation but was shaped by the preceding trial. Whenever participants actively controlled their own search, higher confidence on the previous trial raised confidence on the current one, indicating that subjective certainty propagates across successive judgments even as the underlying evidence changes (49). The two modes of agency differed, however, in the role of prior objective performance. When exploration was screen-based, a correct previous trial further inflated current confidence, whereas when participants explored by moving sensory organs through the environment, current confidence was insensitive to the previous trial’s accuracy. Because the influence of recent history on confidence depends on the sensorimotor context of the decision (50), this pattern suggests that embodied exploration binds confidence more tightly to concurrent sensory input. This behavioral signature aligns with the neural results below, together pointing to a shared embodied grounding that binds confidence to present sensory evidence.

Our study also advances the technical aspects for naturalistic sensory research. SSAEPs index the strength of neural responses to periodic acoustic stimulation (37, 51–53) and have been used to probe auditory attention and stream segregation in stationary listeners (36, 54). Mobile recording of steady-state evoked potentials (SSEPs) during real-world movement, however, has so far been demonstrated mainly for vision, where SSVEPs dissociate the neural response of locomotion from those of visual stimulation (51, 52). This gap matters because metacognition research has relied predominantly on static, passive listening (3, 33, 39), and the few studies linking action to confidence have addressed visual or motor domains rather than naturalistic listening (55, 56). We therefore extend SSEPs to the auditory domain, combining active navigation with trial-by-trial confidence monitoring.

Using this novel approach, we found that agency, whether embodied or screen-based, produced a non-selective increase in SSAEP amplitude, indicating that this enhancement stems from the authorship over self-directed sampling rather than embodiment. The broad increment likely reflects state-dependent neuromodulatory and motor-sensory influences that optimize neural readout during active agency (27, 57); although locomotion can both enhance and suppress auditory-evoked responses (58), we observed consistent enhancement. Crucially, this enhancement did not come at the expense of stimulus selectivity: the target/non-island contrast was preserved across conditions, so perceptual detection remained intact while the overall signal available to higher-order processing increased.

These agency-driven conditions yielded equivalent behavioral performance and non-selective SSAEP enhancement. The target-evoked response, however, was sensitive to confidence, with high-confidence trials eliciting stronger activity than low-confidence trials, and this difference was reliable only under embodied agency. Thus agency is sufficient to raise overall metacognitive sensitivity, whereas embodiment additionally renders the target-evoked response sensitive to subjective certainty. This sensitivity was confined to the target and absent for non-island sounds. Unlike non-target stimuli, targets engage the decisional stage of the cycle (59), where accurate confidence carries the greatest functional weight (60). The target specificity of the effect therefore indicates that embodied agency does not indiscriminately enhance metacognitive readout but selectively strengthens the correspondence between task-relevant neural representations and confidence, consistent with metacognition operating on the decisional evidence accumulated for the task at hand (2, 44). Fluctuations in target-evoked responses may thus provide a neural marker of metacognitive calibration.

Both agency conditions required voluntary action, yet only embodiment physically displaced the sensory organs through space. Embodiment introduces a continuous, body-anchored sensorimotor flow (i.e. vestibular, proprioceptive, and somatosensory signals synchronized with self-generated movement) that screen-based search cannot reproduce (15, 42, 61). Within embodied-cognition frameworks, this multimodal flow does not merely accompany perception but constitutes it, acting as a grounded temporal scaffold that modulates the SSAEP amplitudes and sharpens the temporal sampling on which auditory decisions depend (26, 29, 62, 63). In freely moving animals, for example, auditory cortical neurons adjust their spatial tuning to a sound source’s behavioral relevance during self-motion (64), illustrating the heightened ecological resonance that characterizes naturalistic behavior (28), and the brain–body–environment system implements both perception and its evaluation (65). Because sounds were delivered via headphones without acoustic intensity or spatial gradients, this embodied advantage cannot stem from acoustic changes at the ear, but reflects the locomotor reafference of active exploration itself. Crucially, self-evaluation in this paradigm is not a detached post-decision judgment; it is an online, dynamically ongoing process unfolding during active search (47), which explains why cortical responses (SSAEPs) recorded during ongoing exploration already track the eventual confidence report. By focusing cognitive processing strictly onto concurrent sensory input, embodiment supplies body-anchored precision estimates (66–70) that insulate the observer from prior-trial hysteresis. This real-time grounding seamlessly binds perception, neural representation, and self-evaluation into a complete closed-loop system.

If embodiment tethers self-evaluation to immediate sensory reality, the same principle may extend to current agentic AI architectures, whose self-monitoring remains unreliable: state-of-the-art models are frequently miscalibrated and accurately quantifying their uncertainty is still unsolved (71), with a persistent gap between what a model internally represents and the confidence it expresses (72). Our results suggest that agency alone is insufficient to ground self-monitoring. Embodiment is essential: by anchoring confidence to current sensory evidence and filtering prior history, sensorimotor interaction, rather than virtual control, is key to building agentic AI that monitors its perception reliably.

In conclusion, mobile SSAEPs proved a reliable readout of neural response during naturalistic auditory search, dissociating the contributions of embodiment from agency. While agency was the primary driver of the perceptual enhancement and of the increase in metacognitive sensitivity, embodiment additionally aligned both the neural response and confidence with concurrent sensory evidence. Under embodied search alone, target-evoked neural responses varied with confidence and confidence was insulated from preceding trial history. These findings carry implications for information processing in humans embedded in increasingly screen-based environments, as well as for the development of accurate self-monitoring artificial agents.

## Materials and Methods

### Participants

Twenty healthy adults (11 females; age: *M* = 24.6, *SD* = 3.12) participated. Sample size was determined through G*Power 3.1 (73) to achieve 80% power for a medium effect size (Cohen’s f = 0.25) in a repeated-measures ANOVA. Participants were recruited from LMU Munich via posters and met the inclusion criteria of being right-handed, having normal or corrected-to-normal vision and hearing, and reporting no history of neurological or psychiatric disorders. The experimental protocol was approved by the Ethics Committee of the Faculty of Psychology and Educational Sciences at LMU Munich, with all participants providing written informed consent in accordance with the Declaration of Helsinki. Upon completion of the session, they received monetary compensation or course credit for participation.

### Experimental Design and Procedure

Based on the SITh paradigm (31) structure, we developed a multi-island SITh task. This task included 4 islands, one target (500 Hz) and three non-target islands, each at a different frequency drawn from the set 460, 480, 520, and 540 Hz — three of the four frequencies on any given trial, with the combination varying across trials. A non-island sound (850 Hz) was played outside all the islands. Participants navigated an arena to locate an invisible auditory target “island” hidden among other invisible auditory non-target “islands”, while sounds were delivered binaurally via headphones throughout the experiment (Fig. 1A). There were three conditions: 1) Embodied Agency (EA), embodied exploration in a 3 *×* 3 m arena, the sampled location was reached by displacing the body itself; 2) Stationary Agency (SA), cursor-based navigation on a monitor, in which the sampling was exercised by a hand movement while the body and head remained stationary; 3) NA, a control condition, in which participants passively heard a playback of the exact auditory sequences they had previously generated during their own SA exploration trial by trial (Fig. 1C). Each trial began with three repetitions of the target sound. Following these initial cues, participants began the search phase at their own pace, either by navigating the environment (SA, EA) or by passively listening to the sound stream (NA) to identify the target. What the participant would hear is based on their location in real time (EA & SA). When participants felt they stepped inside the “target” island and heard the “target” stimulus (EA & SA) or purely heard the “target” sound (NA), participants terminated the trial by pressing a button and subsequently reported their confidence on a scale from 1 (completely uncertain) to 10 (completely certain) (Fig. 1B). Each condition comprised 60 trials (each lasting up to 90 seconds). In every agency driven trial (EA and SA), four sound islands were randomly distributed to ensure that performance was driven purely by auditory cues rather than spatial learning or spatial preference (SI Appendix, Figs. S1 and S2). Confidence was split at each participant’s own median, computed separately per condition: trials below that median were labeled low-confidence and the rest high-confidence. The same median-split criterion was applied to every participant and condition (the threshold value itself varied by participant). After finishing the task, participants were asked to complete the Musical Ear Test (MET) (74, 75) as a measure of auditory proficiency. The test consists of 104 trials (52 melodic, 52 rhythmic) that assess melodic and rhythmic discrimination.

The acoustic environment consisted of three kinds of sound: target (50 ms of 500 Hz pure tone sound pips); non-islands (50 ms of 850 Hz pure tone sound pips), and non-target island sounds (50 ms pure tone sound pips of 460, 480, 520, or 540 Hz). These four frequencies lie within the 500 Hz critical band (*∼*79 Hz equivalent rectangular bandwidth); on each trial, three of the four were assigned to the three non-target islands (one frequency per island), creating graded levels of peripheral competition within a single auditory filter. Conversely, the 850 Hz non-island sound was separated from the target by *∼*9 semitones to ensure its robust perceptual segregation from the target and non-target complex (76). Each stimulus lasted 50 ms and was synthesized using a sine-wave carrier with integrated fade-in and fade-out envelopes to prevent acoustic transients and minimize spectral splatter. All auditory stimuli were delivered at a fixed rate of 4 Hz (each sound stimulus consists of 50 ms sound playing and 200 ms blank; each second has 4 sound stimuli), aligned with the original SITh paradigm (30, 31). Beyond paradigm consistency, 4 Hz is suited for eliciting cortically generated SSAEPs: low-rate SSAEPs originate primarily from cortical sources (37, 77), and this frequency aligns with the theta-band range at which the auditory cortex preferentially entrains to rhythmic stimulation (78, 79).

To quantify participants’ metacognitive sensitivity, we computed the area under the type-2 receiver operating characteristic curve (AUROC2), a non-parametric, threshold-free measure (39, 40) that evaluates how well participants’ confidence matches their performance. For each participant and condition, trials were first sorted by accuracy and then, within each accuracy category, by confidence rating. The type-2 ROC curve was constructed by treating each confidence level *c* on the 1–10 scale as a decision criterion and computing, at each criterion, the type-2 hit rate and type-2 false-alarm rate (Eq. 1).

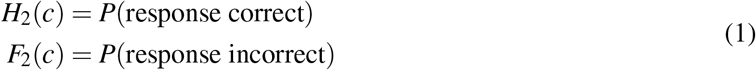

Then plotting *H*_2_(*c*) against *F*_2_(*c*) across all criteria *c* traces the type-2 ROC curve (40), and AUROC2 was computed as the area beneath this curve via trapezoidal integration across successive criteria (Eq. 2).

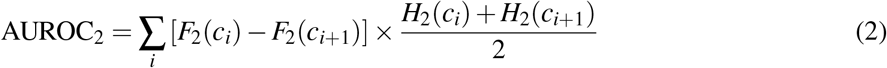

AUROC2 = 0.5 indicates that confidence carries no information about the accuracy of a decision (chance-level metacognitive sensitivity), whereas AUROC2 = 1 indicates that confidence ratings perfectly discriminate correct from incorrect responses. Figure 3D shows the type-2 ROC curves for a representative participant across conditions (EA, SA, NA); the area beneath each curve corresponds to the AUROC2. Additionally, we prioritized AUROC2 to quantify metacognitive sensitivity. As a non-parametric, distribution-free metric, AUROC2 bypasses the restrictive Gaussian assumptions inherent in parametric models, offering a more direct estimate of sensitivity compared with parametric indices such as meta-d’ or meta-d’/d’ (80).

### EEG Recording and Processing

EEG data was recorded with a wireless LiveAmp amplifier (Brain Products GmbH, Gilching, Germany) at a sampling rate of 1000 Hz. The 8-channel montage comprised four scalp electrodes (Cz, Fz, T7, T8), two earlobe electrodes, and two electro-oculogram (EOG) channels for monitoring eye movements, together with dedicated online reference (FCz) and ground (AFz) electrodes. All analyses reported in the main text use the frontal midline electrode Fz. SSAEPs are most commonly recorded from a single channel between a midfrontal or vertex electrode, and are largest over frontocentral scalp regions (37). Waveforms at the remaining recorded channels (Cz, T7, T8) are shown in SI Appendix, Fig. S3. Electrode impedances were maintained below 5 kΩ throughout recording. All subsequent processing was performed offline in MNE-Python (81). Data were first re-referenced to the average of the two earlobe electrodes. For analyses based on the stimulus-locked evoked waveform, the re-referenced signal was band-pass filtered between 3 and 20 Hz and segmented into 250-ms epochs time-locked to each auditory stimulus onset. Epochs in which the signal exceeded *±*100 µV at any channel were excluded from further analysis (mean *±* SD rejection rate: 1.9 *±* 5.1% in EA, 0.0 *±* 0.1% in SA, and 0.1 *±* 0.1% in NA; no participant lost more than 25% of epochs in any condition, maximum = 23.3%), yielding comparably clean evoked-response estimates across the three conditions.

For the frequency-domain characterization of SSAEPs (Fig. 2A), a parallel preprocessing branch was used. The re-referenced EEG was segmented into stimulus-driven epochs (within trials) and non-stimulus-driven epochs (during inter-trial intervals) without filtering. Stimulus-driven and non-stimulus-driven segments were concatenated across runs within each condition (EA, SA, NA) for each participant, and the PSD was estimated using Welch’s method (20-s window length, 50% overlap, over 0.01 – 45 Hz). To express spectral power at each frequency relative to the local noise floor, we computed a normalized PSD, following the approach established for steady-state evoked potentials by Meigen & Bach (82) and implemented analogously in MNE-Python’s SSVEP analysis pipeline. For each frequency bin *f*, the PSD was divided by the mean PSD of the twelve neighboring bins (six on either side of *f*, spanning approximately *±*0.1–0.4 Hz), excluding the bin immediately adjacent to *f* on each side so that the local baseline estimate is not contaminated by spectral leakage from *f* itself (Eq. 3).

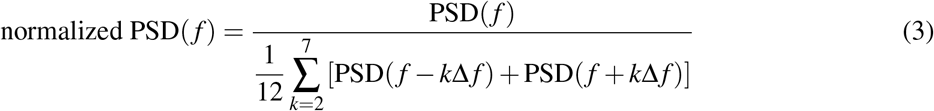

Values near 1 indicate spectral power indistinguishable from the local noise floor, whereas values exceeding 1 reflect frequency-specific neural entrainment. This normalization was applied separately for each participant, condition, and segment type, producing the sharp, comparably scaled peaks at 4 Hz and its harmonics (8, 12, 16, and 20 Hz) during stimulus-driven epochs relative to the flat spectral profile during non-stimulus-driven epochs (Fig. 2A).

As a complementary time-domain assessment of signal stability, we evaluated the test-retest reliability of the SSAEP waveform evoked by the non-island stimulus. For each participant and condition, all 250-ms stimulus-locked epochs corresponding to the non-island stimulus were extracted from the preprocessed data described above and divided into two halves according to trial order (first half, second half). Each half was averaged separately to obtain two waveforms, and the Pearson correlation between these two averaged waveforms at electrode Fz was taken as the reliability coefficient for that participant and condition (Fig. 2B; check supplementary for other channels). This split-half procedure estimates the consistency of the stimulus-evoked response across the recording session, independently of its absolute amplitude.

Following the procedures described by Dowsett (52, 53), the strength of the stimulus-locked SSAEP was quantified as the peak-to-trough (PTT) amplitude of the averaged waveform of sound stimuli played at a rate of 4 Hz, computed separately for target and non-island stimuli. To prevent unequal epoch counts across conditions and stimuli from biasing the evoked responses, PTT amplitude was estimated by bootstrap. Within each participant and cell, epochs were resampled with replacement to that participant’s smallest cell size (10,000 iterations); PTT was computed on the averaged waveform of each iteration, and the 10,000 resulting values were averaged to yield the participant’s PTT for that cell. This time-domain approach has the advantage of accommodating non-sinusoidal waveform shapes without requiring a priori selection of which harmonics to include, a limitation inherent to frequency-domain and spectral power methods. For an averaged waveform *V* (*t*) over the epoch window, PTT was defined as the difference between its maximum and minimum amplitude (Eq. 4).

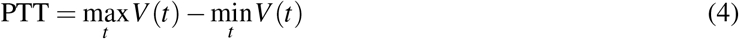

To constrain the influence of broadband noise and movement-related artifacts, we additionally extracted control epochs using a “fake” stimulation rate of 3.1 Hz in place of the true 4 Hz sounds playing trigger rate, applying the same 250-ms epoch duration and the same within-trial extraction procedure as the real stimulus-locked epochs. Because 3.1 Hz is non-harmonic and not locked to the stimulus train, the PTT amplitude of the resulting averaged waveform provides an estimate of the local noise floor against which the genuine stimulus-evoked response can be referenced. The normalized amplitude, which shares the same calculation logic as signal-to-noise ratio (SNR), was then defined as the ratio of the stimulus-locked PTT at 4 Hz to the control PTT at 3.1 Hz (Eq. 5).

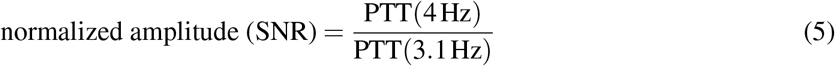

### Statistics

Behavioral condition effects were assessed using repeated-measures ANOVA; where distributional assumptions were not met, Friedman tests were used as the nonparametric alternative. As SSAEP amplitudes were non-normal (Shapiro–Wilk), factorial effects used ART-ANOVA with marginal-means post-hoc comparisons; within-condition contrasts (high-vs low-confidence, correct vs incorrect) used Wilcoxon signed-rank tests. The significance threshold was set at *α* = .05 throughout. Where multiple comparisons were performed, false discovery rate (FDR) correction was applied within each comparison family unless otherwise stated.

## Data, Materials, and Software Availability

All behavioral data, preprocessed mobile EEG datasets, and custom analysis scripts will be deposited in OSF and will be publicly available upon publication. During peer review, data and code are available from the corresponding authors Michael Pecka and Dardo N. Ferreiro upon reasonable request.

## Acknowledgments

We thank Valerie Herbert for assistance with participant recruitment and EEG recording, and Sven Schoernich for providing technical solutions. We thank Bahador Bahrami and Ophelia Deroy for their inspiration in investigating metacognition using the SITh paradigm. This work was supported by DFG grant PE2251/4-1 to M.P. and DO 2460/1-1 to J.D.; Z.T. was supported by the LMU-CSC Scholarship. EEG data were collected at LMU Biocenter. Use of Claude (Anthropic, Claude Sonnet 4.6) for editorial assistance in manuscript preparation is disclosed in accordance with journal policy.

## Author Contributions

**Z.T**.: Methodology, Software, Formal Analysis, Investigation, Data Curation, Writing – Original Draft, Visualization. **D.N.F**.: Conceptualization, Supervision, Writing – Review & Editing. **M.P**.: Conceptualization, Resources, Writing – Review & Editing, Supervision, Funding Acquisition. **P.T**.: Methodology, Supervision, Writing – Review & Editing, Funding Acquisition. **J.D**.: Methodology, Writing – Review & Editing, Funding Acquisition.

