## Supplementary for "Agency and Embodiment Provide Dissociable Benefits to Perception, Metacognition and Neural Processing During Active Sensing"

**This PDF file includes:**

Figures S1 to S8

### Supplementary Figures

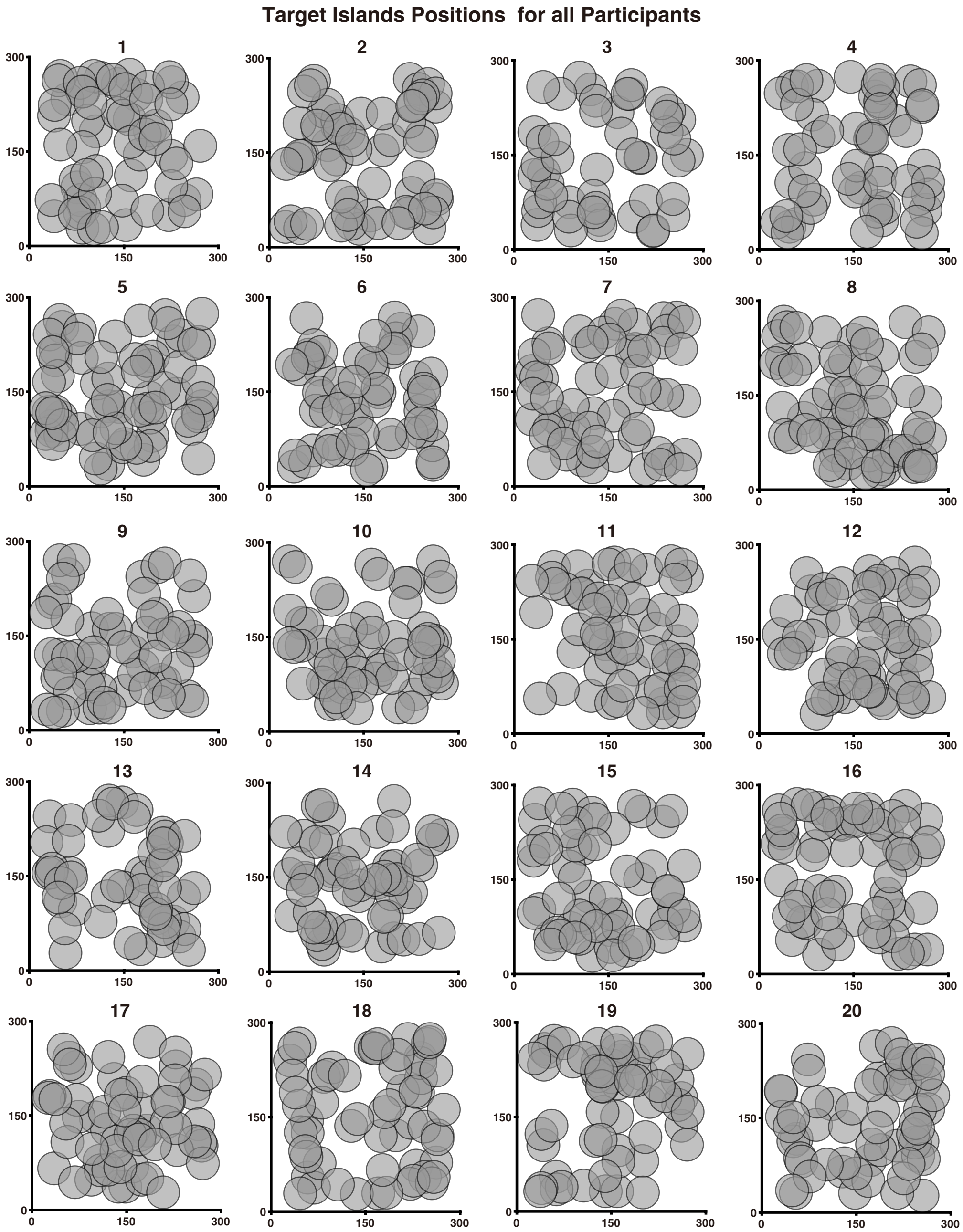


**Fig. S1. Spatial Distribution of Target Island Positions Across Participants.** Locations of the target island pooled across trials in the active conditions (EA and SA). Each subplot shows one session of one participant; target positions are distributed approximately uniformly across the arena, confirming that successful search was driven by auditory cues rather than spatial learning or location bias.

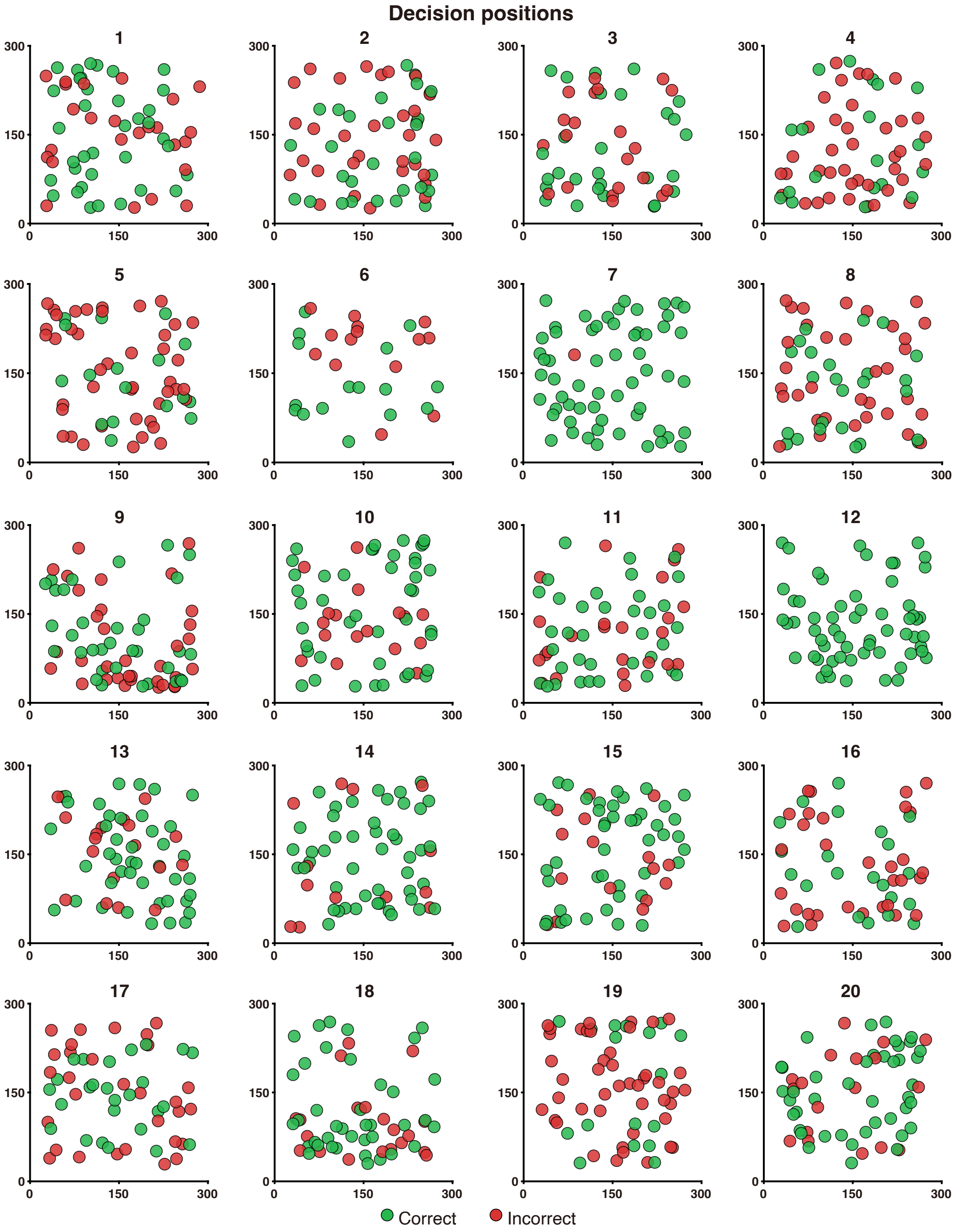


**Fig. S2. Spatial Distribution of Decision Positions Across Participants.** Locations at which participants terminated their search (decision positions) pooled across trials in the active conditions (EA and SA), colored by outcome (green, correct; red, incorrect). Each subplot shows one session of one participant; decision positions are distributed approximately uniformly across the arena, indicating no systematic spatial bias in where participants chose to respond.

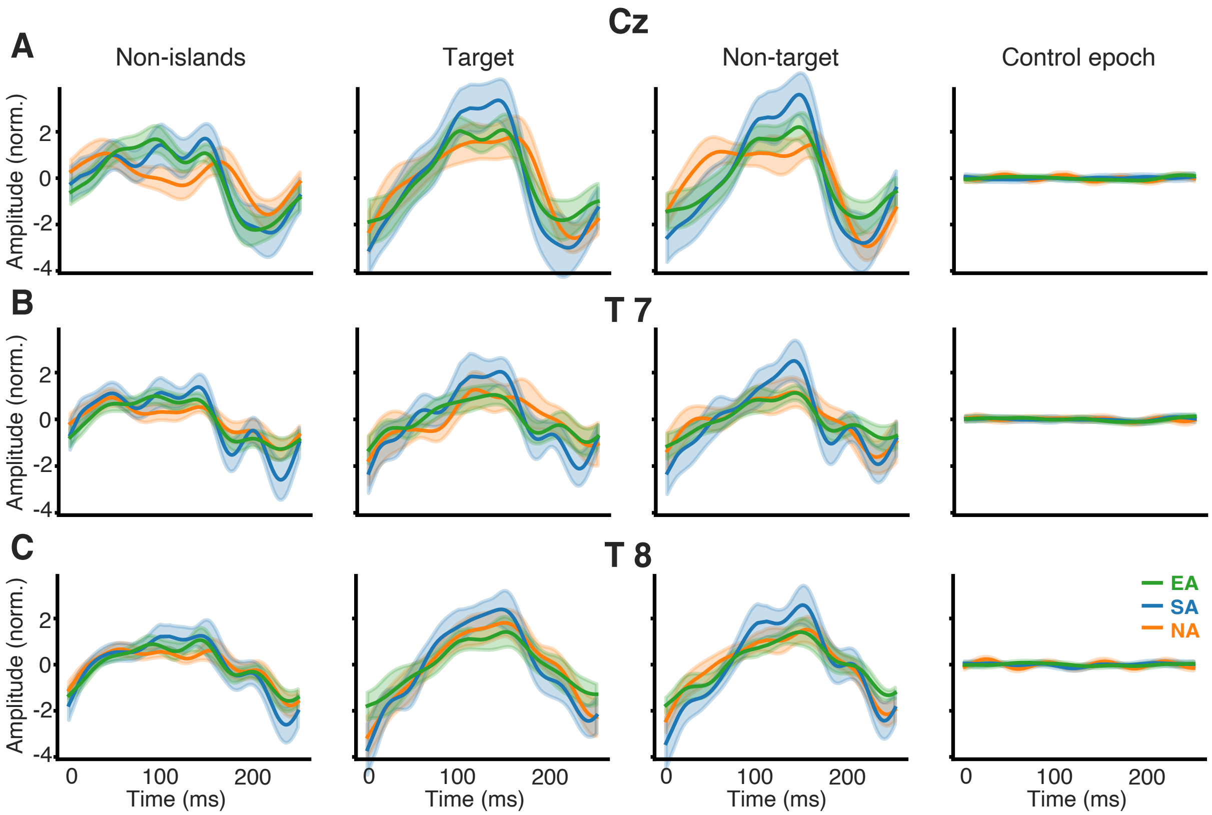


**Fig. S3. Evoked Waveforms across Sound Stimuli and Channels.** Normalized waveforms of Cz (**A**), T7 (**B**), and T8 (**C**). Time-domain SSAEP waveforms for non-island, target, and non-target island sounds exhibit distinct, component-specific waveforms, in contrast to the near-flat response obtained from control epochs analyzed at the 3.1 Hz non-stimulus frequency. Waveforms span a full 250 ms cycle (0–250 ms) with shaded error bands.

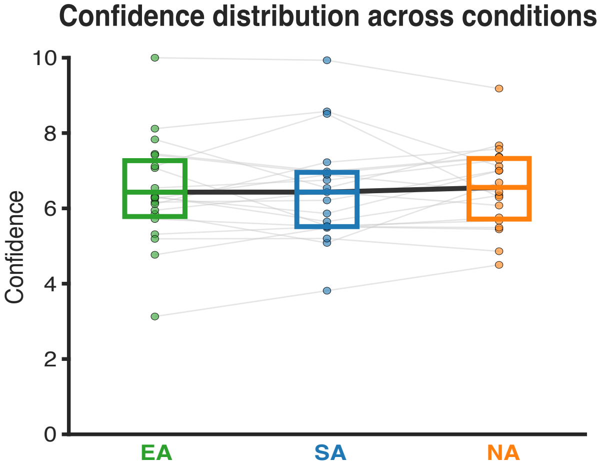


**Fig. S4. Confidence Ratings Across Conditions.** Mean subjective confidence rating per participant in each condition (EA, SA, NA), averaged across all trials within a condition. Confidence did not differ significantly across the three conditions (repeated-measures ANOVA, $F(2,38) = 0.32$, $p = .73$).

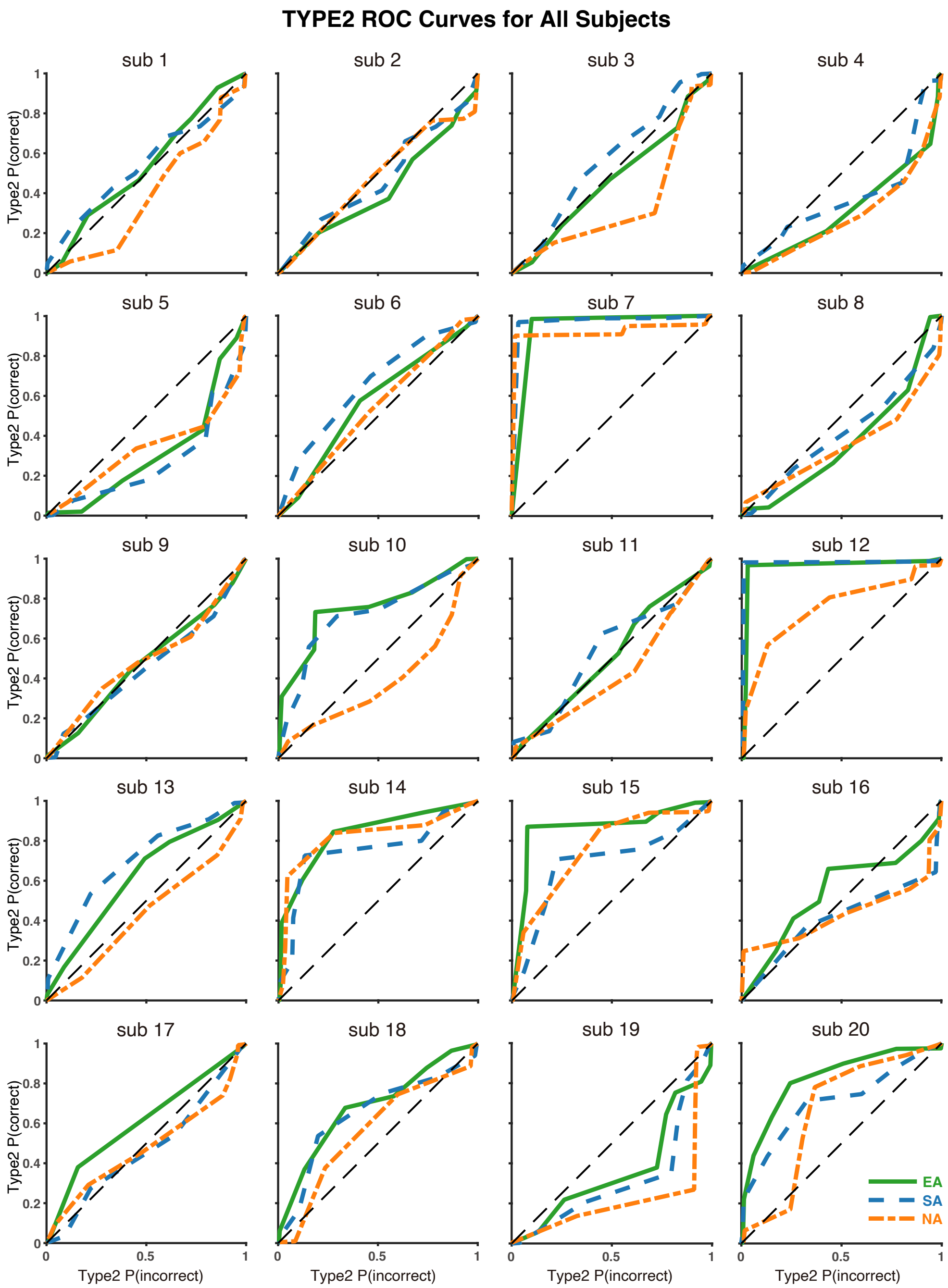


**Fig. S5. Type-2 ROC Curves for All Participants Across Conditions.** Individual type-2 ROC curves for every participant in each of the three conditions (EA, SA, NA), showing how subjective confidence tracks objective performance; the area under each curve is that participant's $AUROC_2$. Figure 3D shows the curves of participant 10.

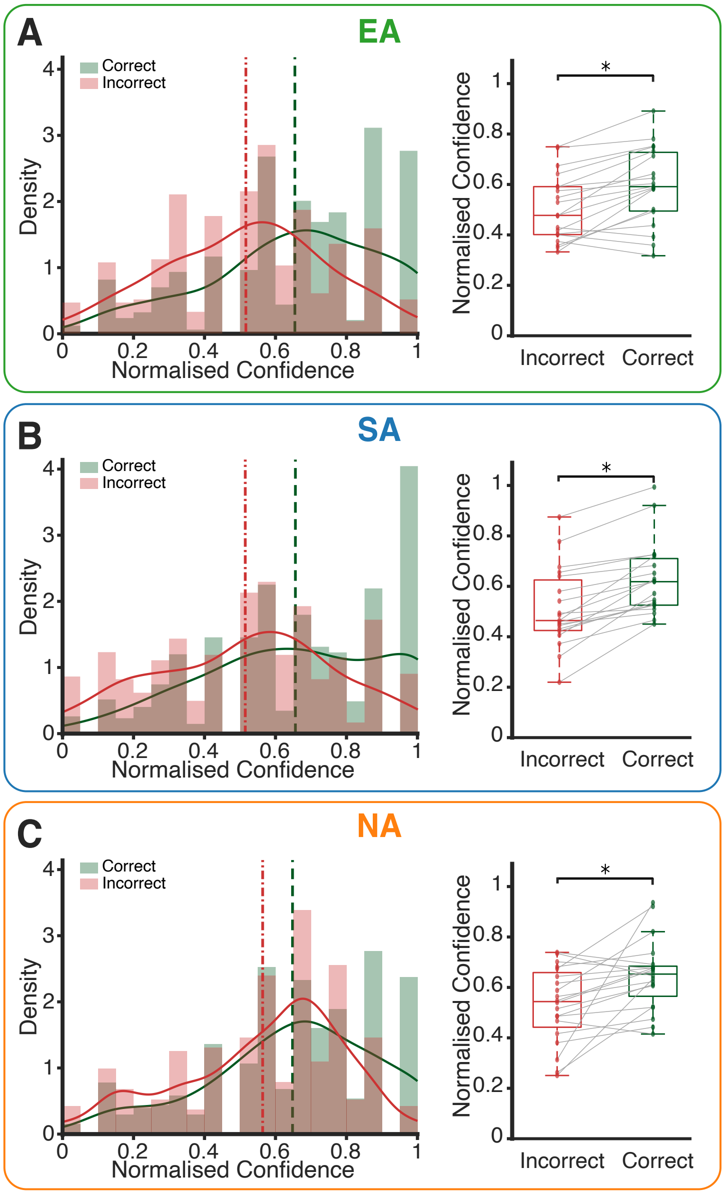


**Fig. S6. Confidence Is Higher on Correct than Incorrect Trials in All Conditions.** Normalized confidence distributions for correct versus incorrect trials, shown separately for each condition; within-condition comparisons used Wilcoxon signed-rank tests. (**A**) EA: confidence was significantly higher on correct than incorrect trials (N = 19, p = .002). (**B**) SA: the same pattern (N = 19, p < .001). (**C**) NA: the same pattern (N = 20, p = .002). Across all conditions, higher confidence accompanied correct target identification, confirming that subjective confidence tracked objective accuracy.

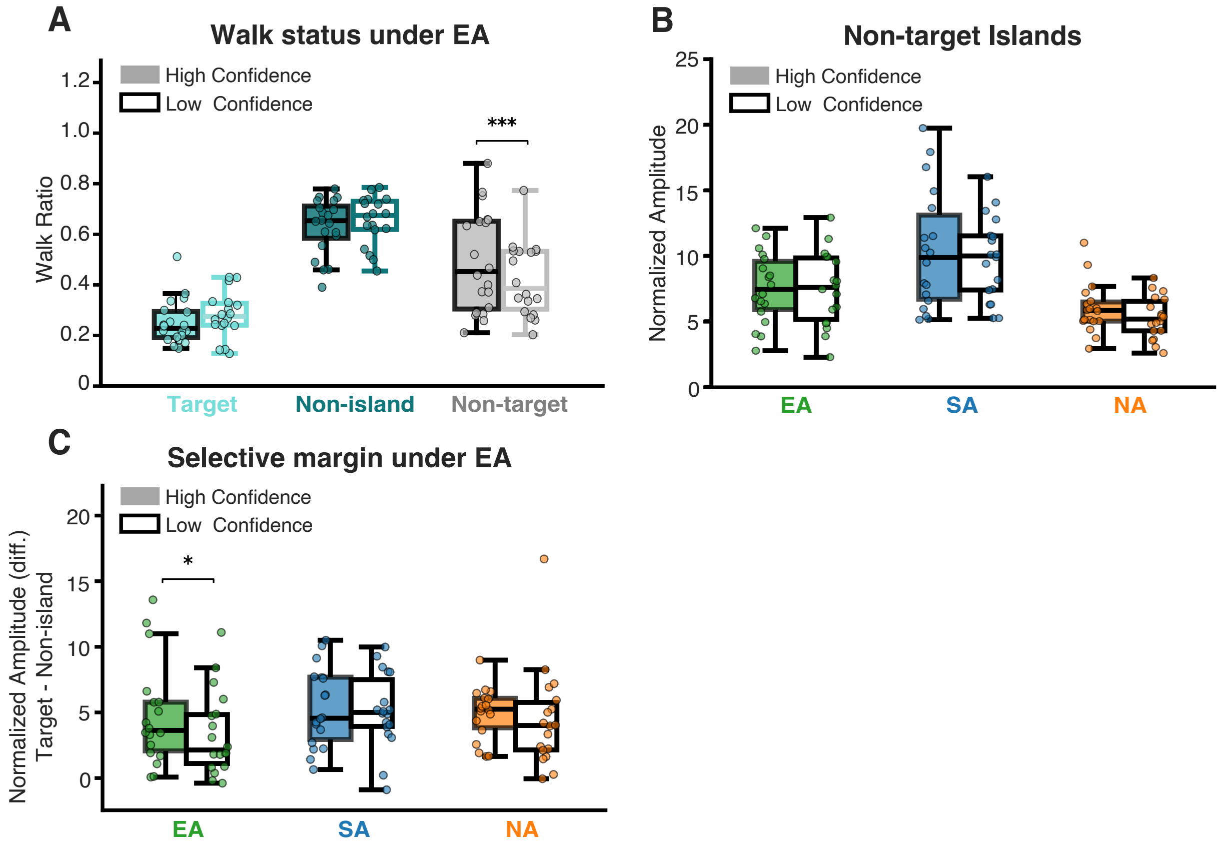


**Fig. S7. Confidence Modulation Is Specific to the Target and Not Driven by Walking.** (**A**) *Walking ratio by stimulus type and confidence during EA.* Fraction of time spent walking during stimulus presentation for target, non-island, and non-target-island stimuli, split by confidence state (N = 18, paired t-tests). Walking did not differ between high- and low-confidence trials for the target (p = .35) or the non-island stimulus (p = .99), and was higher on high-confidence trials only for the non-target islands (p < .001). (**B**) *Stability of non-target-island processing across confidence levels at Fz.* Normalized SSAEP amplitude evoked by the non-target islands was not modulated by confidence in any condition (3 × 2 repeated-measures ANOVA: confidence main effect F(1,17) = 2.85, p = .11; condition × confidence interaction F(2,34) = 0.76, p = .48; N = 18; within-condition high-vs-low post hoc all $p_{corr} \geq .08$, FDR-corrected). The condition main effect (F(2,34) = 17.89, p < .001) reflects the overall amplitude increase during active search. This parallels the absence of confidence modulation for the non-island stimulus (Fig. 5E). Participants 7 and 17 are excluded for having no low-confidence trials. Because non-target-island amplitude nonetheless showed no confidence modulation (**B**), the walking difference in (**A**) did not produce an amplitude difference, confirming that movement is removed by preprocessing and does not shift the neural response across confidence states. Consequently, the higher target-evoked SSAEP amplitude on high-confidence trials during EA (Fig. 5D) cannot be attributed to greater movement. (**C**) *Selectivity margin by confidence state.* Normalized amplitude difference (target minus non-island) at Fz, split by confidence state in each condition. The margin was larger on high- than on low-confidence trials only under EA (t(17) = 2.38, p = .03, d = 0.56), and not under SA (p = .72) or NA (p = .54); this is the boxplot rendering of the data shown in Fig. 5F. Boxplots display medians and quartiles with individual subject data points.

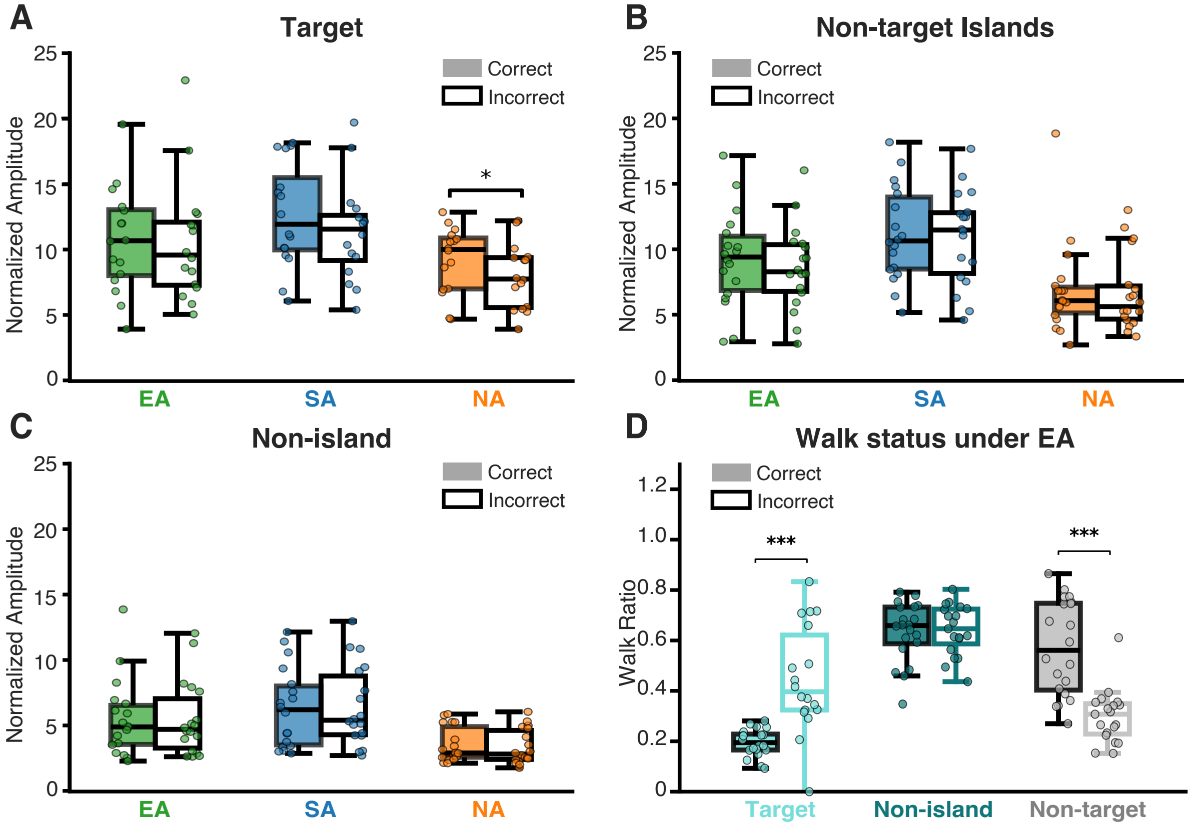


**Fig. S8. SSAEP Amplitude at Fz and Trial Accuracy.** (**A–C**) *SSAEP amplitude by accuracy.* Normalized amplitude for the target (**A**), non-target islands (**B**), and non-island (**C**) stimuli at Fz, split by trial accuracy (correct vs incorrect) within each condition. For the target (**A**), a 3 × 2 repeated-measures ANOVA revealed a main effect of accuracy ($F(1,15) = 9.21$, $p = .008$) and of condition ($F(2,30) = 6.65$, $p = .004$), with no condition × accuracy interaction ($F(2,30) = 1.03$, $p = .37$; $N = 16$); post hoc tests localized the accuracy effect to passive listening (correct > incorrect, $p < .001$, FDR-corrected), with no difference during EA ($p = .78$) or SA ($p = .19$). Non-target island amplitude (**B**) showed no accuracy modulation (accuracy $F(1,17) = 1.53$, $p = .23$; interaction $F(2,34) = 0.28$, $p = .75$; $N = 18$). Non-island amplitude (**C**) showed no accuracy modulation (accuracy $F(1,17) = 0.01$, $p = .93$; interaction $F(2,34) = 0.54$, $p = .59$; $N = 18$). (**D**) *Walking ratio by stimulus type and accuracy during EA.* Fraction of time spent walking during stimulus presentation for target, non-island, and non-target-island stimuli, split by accuracy ($N = 19$; Wilcoxon signed-rank). Walking did not differ between correct and incorrect trials for the non-island stimulus ($p = .97$) but differed in mirror-image fashion for the target (correct < incorrect, $p < .001$) and the non-target islands (correct > incorrect, $p < .001$), reflecting where the search terminated: on correct trials participants stopped inside the target island (less walking during target sounds, more during non-target sounds), whereas on incorrect trials they stopped inside a non-target island (the reverse). Boxplots display medians and quartiles with individual subject data points.
